## Supplementary Materials for "Quantification of *Myxococcus xanthus* Aggregation and Rippling Behaviors: Deep-learning Transformation of Phase-contrast into Fluorescence Microscopy Images"

### 1. Text S1: Supplemental Methods

#### *1.1. Image processing*

##### 1.1.1. Downscaling

The raw 2048 × 2044 images are encoded in 2-byte (16-bit) unsigned integers. Since lower resolution images are good enough for cell density estimation, the 0.65 micros/pixel images were 2× downsampled to 1.30 micros/pixel by LANCZOS resampling algorithm [1]. This also reduces storage memory requirements. The LANCZOS resampling uses a convolution kernel to interpolate the pixels of the input image. The kernel function is defined as:

$$k(x) = \begin{cases} \operatorname{sinc}(x)\operatorname{sinc}(\frac{x}{a}) & -a < x < a \\ 0 & \text{otherwise} \end{cases} \tag{1}$$

Here, $x$ is the position relative to the kernel center and $a$ is the kernel size. We used the default settings in python-pillow [2]. The LANCZOS resampling reduces aliasing while maintaining the sharpness of downsampled images [1].

##### 1.1.2. Normalization

To make the images compatible with our model, we normalized the image intensity to $[-1, 1]$ float32 values. For the tdTomato fluorescent images, we applied histogram equalization (HE) [3] to visualize the detailed information, such as the cell streams between different aggregates. The new intensity $I_{new}$ is calculated by the following equation:

$$I_{new} = \frac{I_{raw} - I_{min}}{I_{max} - I_{min}} \times 2 - 1 \tag{2}$$

where $I_{raw}$ is the raw image intensity. $I_{max}$ and $I_{min}$ are the maximum and minimum respectively of the raw image intensity. Due to the differences in the light source, some parts of the images may be brighter than the other parts. To reduce the imbalance in the light condition, we applied contrast limited adaptive histogram equalization (CLAHE). The CLAHE utilizes the pixel intensity distribution from a neighborhood region, which is defined by a tile grid size, and clips the histogram at a predefined value, called the clip limit. We set the clip limit equal to 2, and the tile grid size to 16 by 16 pixels.

##### 1.1.3. Augmentation

We have collected 9 movies of $5 \times 10^9$ cells/mL, 3 movies of $2.5 \times 10^9$ cells/mL and 2 movies of $1 \times 10^{10}$ cells/mL each containing 1440 or more frames (24hr of images taken every minute). For the training set, we picked images in 7 randomly selected movies of $5 \times 10^9$ cells/mL and 3 randomly selected movies of $2.5 \times 10^9$ cells/mL at the interval of 10 minutes. We then manually removed the images captured when the light bulb was off. Images from one of the $5 \times 10^9$ cells/mL movies form the cross-validation set, and images from the other 2 movies at the same density are annotated as the test set. The model is trained on the training set and the model performance is evaluated on all the images in the 12 movies. The training set should be large enough to contain images of all possible strains, so that our model can learn to generate tdTomato fluorescent images from different experimental phase-contrast images. To increase the variety of images in the training set, we randomly selected positions on the image, and cropped the images to 256×256 pixels, which is large enough to contain at least one aggregate on average in the image, and small enough to be processed in our model. Then we randomly rotated the downscaled input and ground truth images together by 0°, 90°, 180° or 270°. Finally, we randomly mirrored 50% of the images.

Data augmentation is important for such tasks with limited datasets. For a 1024×1024 pixel image, there are as many as $4(crop) \times 4(crop) \times 4(rotate) \times 2(flip) = 128$ different small 256×256 pixel images. These images represent the same time frame of an experiment but have aggregates distributed differently. Without data augmentation, it is possible to see an aggregate existing on the same corner across different images. Our model may overfit and treat that fixed aggregate as an underlying feature of the fluorescent image. The data augmentation leads to variations in aggregate distribution and helps our model learn the image transformation regardless of the image orientation and distribution of aggregates.

##### 1.1.4. Image Segmentation

The aggregates are sized between a lower bound and an upper bound. To get the best lower bound and upper bound for segmentation, we started from the aggregate size range determined in [4], where the lower and upper bounds are 7 and 1000 pixels in our image. However, it takes longer to segment with such a large upper bound. We have tested the aggregate segmentation with smaller upper bounds and found that with 200 pixels as the upper bound, the result does not change too much (Figure S1), but it is about 45 times faster. Therefore, we picked 7 and 200 as the lower and upper bounds.

#### *1.2. Network Architectures*

In this section, we will first introduce the pix2pixHD model, and then the modification in pix2pixHD-HE. Finally, we will discuss the training details.

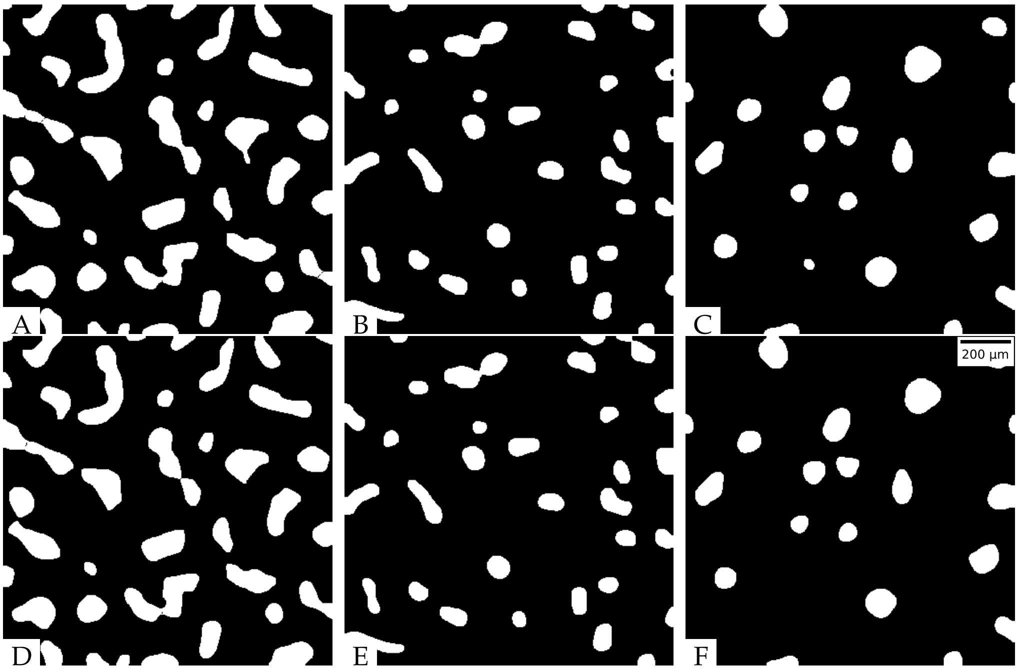

**Figure S1.** Aggregates segmented at the 800 minute, the real fluorescent images are shown in Figure 6C, S9C, S10C. (A)1 $\times$ $10^{10}$ cells/mL, upper bound=200, execution time: 1.3s; (B)5 $\times$ $10^{9}$ cells/mL, upper bound=200, execution time: 1.3s; (C)2.5 $\times$ $10^{9}$ cells/mL, upper bound=200, execution time: 1.4s; (D)1 $\times$ $10^{10}$ cells/mL, upper bound=1000, execution time: 61.5s; (E)5 $\times$ $10^{9}$ cells/mL, upper bound=1000, execution time: 61.9s; (F)2.5 $\times$ $10^{9}$ cells/mL, upper bound=1000, execution time: 61.4s;

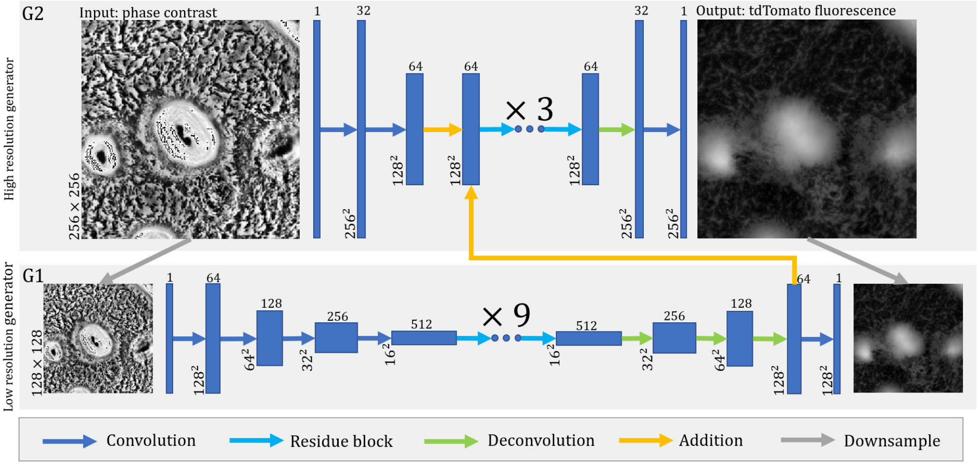

**Figure S2.** The network architecture of pix2pixHD generator. Each rectangular block represents a 3-dimensional array. The lengths and widths of the arrays are annotated on the left. The depths of the arrays are annotated on the top. The blue dots represent several 2-layer residual building blocks. The numbers of blocks are annotated above.

#### 1.2.1. Pix2pixHD Generator

We base our method on the pix2pixHD [5] (Figure S2) model, which is a conditional GAN designed for high-resolution image transformation, to process our images. The model has two levels of generators, $G_1$ and $G_2$. The architectures are shown as follow:

$G_1$: C64–C128–C256–C256–C512–R512–R512–R512–R512–R512–R512–R512–R512–R512–D256–D128–D64–C1

$G_2$: C32–C64–(+$G_1$@D64)–R64–R64–R64–D32–C1

Here $Ck$ denote a Convolution-BatchNorm-ReLU layer with $k$ filters (in the last layers of $G_1$

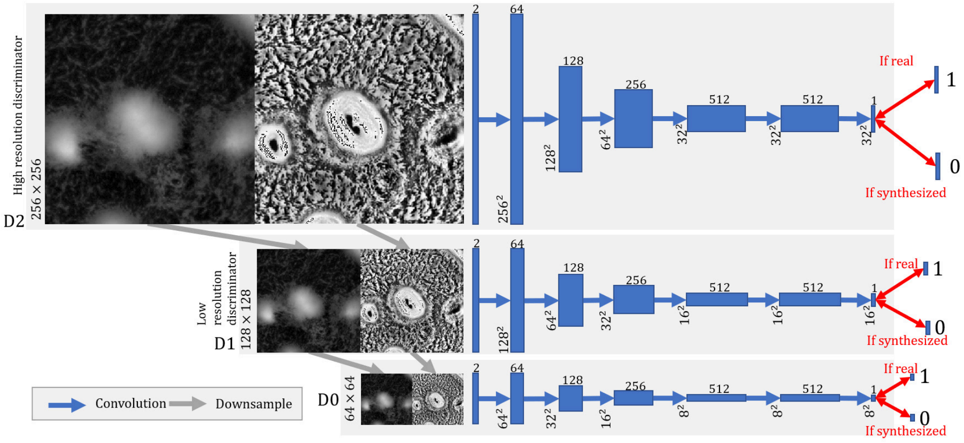

**Figure S3.** The network architecture of multi-scale discriminators. Each rectangular block represents a 3-dimensional array. The lengths and widths of the arrays are annotated on the left. The depths of the arrays are annotated as superscripts. Each arrow represents an operation step. The double-headed arrow represents the MSE calculation.

and $G_2$, the activation function is changed to tanh). R$k$ denotes a Convolution-BatchNorm-ReLU-50%Dropout-Convolution-BatchNorm-Addition residue block[6] with $k$ filters. D$k$ denotes a Deconvolution-BatchNorm-ReLU layer with $k$ filters. Element-wise addition is denoted as $+$, and $G_1$@D64 stands for D64 layer of $G_1$.

#### 1.2.2. Multi-scale Discriminators

We used multi-scale discriminators ($D_0$, $D_1$, $D_2$) to determine whether an image is a real microscopy image (Figure S3). These discriminators follow identical 6-layer convolution network structure[5]:

C64–C128–C256–C512–C512–C1

The activation function is LeakyReLU with 0.2 negative slope. There is no normalization on the first and last layers. There is no activation function on the last layer.

The $D_0$, $D_1$, and $D_2$ were trained to differentiate real and synthesized tdTomato images at 3 different scales. Though taking the same input, the outputs of $D_0$, $D_1$, and $D_2$ are $8 \times 8$, $16 \times 16$, and $32 \times 32$. Thus, elements in the outputs of $D_0$, $D_1$, and $D_2$ have decreasing inspection fields. The $D_0$ focuses on the global information, $D_1$ pays more attention to the mid-level information, and $D_2$ handles the local information. This architecture is compatible with profile and detail transference. During training, the phase-contrast image and tdTomato fluorescent image were stacked together and downscaled by $2 \times 2$ and $4 \times 4$ average polling kernels. Then the original and downscaled image stacks were fed into discriminators $D_2$, $D_1$, and $D_0$. The GAN loss:

$$\min_{G_i} \max_{D_0,\ldots,D_i} \sum_{k=0}^{i} \mathcal{L}_{GAN}(G_i, D_k) \tag{3}$$

where $i = 1$ for training $G_1$, $i = 2$ for training $G_2$. The objective function $\mathcal{L}_{GAN}(G_i, D_k)$ is given by:

$$\mathcal{L}_{GAN}(G_i, D_k) = \mathbb{E}_{(p,t)}[\log D_k(p,t)] + \mathbb{E}_p[\log(1 - D_k(p, G(p)))] \tag{4}$$

where $p$ is the phase-contrast image, $t$ is the fluorescent image. $\mathbb{E}$ represents the expected value.

#### 1.2.3. Feature Matching Loss

Our synthesized fluorescent images should have similar feature distributions as real fluorescent images. Therefore, we minimized the L1-norm between the discriminator outputs of real image stacks and synthesized image stacks on each layer. The feature matching loss is:

$$\mathcal{L}_{FM}(G_i, D_k) = \mathbb{E}_{(p,t)} \sum_{j=1}^{H} \frac{1}{N_j} ||D_k^{(j)}(p,t) - D_k^{(j)}(p, G_i(p))||_1, \tag{5}$$

where $p$ is the phase-contrast image, $t$ is the fluorescent image, $H$ is the total number of layers, and $N_i$ denotes the number of elements in each layer.

The optimization function with discriminator loss and feature matching loss:

$$\min_{G_i} \max_{D_k} \sum_k \mathcal{L}_{GAN}(G_i, D_k) + \lambda \sum_k \mathcal{L}_{FM}(G_i, D_k), \tag{6}$$

where $\lambda$ controls the weight of feature matching loss, and it has a value between 0.1 and 1. During training, we gradually remove the loss function of $G_1$ to avoid over-fitting on profile scale.

##### 1.2.4. Hyperparameter Optimization in Pix2pixHD-HE

We manipulated the model to have two outputs and named it pix2pixHD-HE. The CLAHE output forces our model to learn profile aggregation information, while the HE output assists our model to learn detailed stream information. The pix2pixHD-HE is a variant of pix2pixHD, and the best branch point for the histogram equalized output is unreported in previous publications. To determine the best branch point, we switched the branch point from the second-to-last layer to the first residual block, and labeled them from 1 to 5 (Figure 2). We trained each model for 2000 epochs, and tested them on the cross-validation dataset. The performances were evaluated by MSE and SSIM in Tables S1, S2. In both cases, the branch point at the third to last layer(label 2, Figure 2) performed the best, which we use as our model structure. Using the same annotation as described in Appendix 1.2.1 with the bracket with subscript denotes branched outputs, the architectures are as follows:

$G_1$: C64–C128–C256–C256–C512–R512–R512–R512–R512–R512–R512–R512–R512–R512–D256–D128–D64$(-C1)_2$

$G_2$: C32–C64–(+$G_1$@D64)–R64–R64–R64$(-D32-C1)_2$

Table S1: Hyperparameter optimization, SD: spatial displacement ($\mu m$), $\sigma$: minimal standard deviation for added Gaussian noise, given as a percentage of the original intensity scale, Difference: Subtract the metric value of the best model from the metric value of the current model. The row of the best model used for all the figures is highlighted in bold.

| | MSE | | | | SSIM | | |
|---|---|---|---|---|---|---|---|
| **Branch Point** | **Value** | **SD** | $\sigma$ | **Difference** | **Value** | **SD** | **Difference** |
| 1 | 0.034±0.019 | 7.0 | 9.3% | -0.009±0.010 | 0.663±0.088 | 2.0 | 0.041±0.031 |
| **2** | **0.026±0.011** | **5.0** | **8.0%** | **0.000±0.000** | **0.705±0.073** | **1.9** | **0.000±0.000** |
| 3 | 0.027±0.012 | 5.1 | 8.2% | -0.001±0.002 | 0.694±0.075 | 1.9 | 0.010±0.006 |
| 4 | 0.026±0.011 | 5.2 | 8.1% | -0.001±0.003 | 0.693±0.064 | 2.0 | 0.012±0.017 |
| 5 | 0.033±0.017 | 6.7 | 9.0% | -0.007±0.009 | 0.677±0.080 | 2.0 | 0.028±0.026 |
| None | 0.025±0.010 | 4.9 | 8.0% | 0.001±0.002 | 0.697±0.070 | 2.0 | 0.007±0.008 |

##### 1.2.5. Training Details

All the networks were trained from scratch. We used the Adam optimizer [7] with a learning rate of $5 \times 10^{-5}$ and momentum parameters $\beta_1 = 0.5$, $\beta_2 = 0.999$, the other parameters were set to default ones in pix2pixHD. The model was trained and tested on an HPE XL190 server, with an Intel(R) Xeon(R) Gold 6230 CPU and 32Gb Tesla V100 GPU. For each epoch, the model was trained on 1020 256 × 256 images cropped from 1020 1024 × 1022 images, with 40 to 45 images in each batch. We first trained the $G_1$ and $G_2$ together for 500 epochs, the weight of loss for $G_1$ decreases linearly over time. After

Table S2: Hyperparameter optimization after histogram equalization, SD: spatial displacement ($\mu m$), $\sigma$: minimal standard deviation for added Gaussian noise, given as a percentage of original the intensity scale, Difference: Subtract the metric value of the best model from the metric value of the current model. The row of the best model used for all the figures is highlighted in bold.

| | MSE | | | | SSIM | | |
|---|---|---|---|---|---|---|---|
| **Branch Point** | **Value** | **SD** | $\sigma$ | **Difference** | **Value** | **SD** | **Difference** |
| 1 | 0.177±0.042 | 3.4 | 21.1% | -0.023±0.015 | 0.337±0.070 | 2.6 | 0.051±0.028 |
| **2** | **0.154±0.034** | **3.0** | **19.6%** | **0.000±0.000** | **0.388±0.069** | **2.3** | **0.000±0.000** |
| 3 | 0.167±0.031 | 3.3 | 20.4% | -0.012±0.007 | 0.364±0.059 | 2.4 | 0.024±0.015 |
| 4 | 0.167±0.034 | 3.2 | 20.4% | -0.013±0.011 | 0.356±0.063 | 2.4 | 0.032±0.025 |
| 5 | 0.181±0.039 | 3.5 | 21.2% | -0.026±0.016 | 0.333±0.068 | 2.6 | 0.055±0.031 |
| None | 0.162±0.032 | 3.1 | 20.1% | -0.008±0.009 | 0.369±0.063 | 2.4 | 0.020±0.020 |

500 epochs, we stopped the training of $G_1$ to avoid overfitting downscaled images and to save training time. The model can also process any multiple of $16\times16$ images, such as $1024 \times 1024$ or $768 \times 256$ images.

##### 1.2.6. Model Building Interface

Our models are built in .json files and processed with PyTorch [8]. To make the model building process simpler, we have also made a model builder interface with React and host it on surge (http://igoshinlab-mbuilder.surge.sh/). The website code is also available on GitHub (https://github.com/IgoshinLab/ModuleBuilder.git). We can import a model on the website and edit each layer. The right side of the webpage shows the building blocks. Different layer types can be selected from the drop-down list. The parameters of the selected layer type are specified below that. After pressing the "Submit" button, we can see the corresponding layer added or edited on the left. After we have completed building the model, we can export it locally.

#### *1.3. Mean Square Error and Structural Similarity Indexed Measure Analysis*

##### 1.3.1. Spacial Displacement

For a given initial image position $(\rho, \theta)$ in polar coordinates for a real image $r$ and a generated image $g$, there exists a smallest radial perturbation $\Delta\rho$, such that for all $\theta$

$$MSE(r(\rho,\theta), g(\rho,\theta)) \leq MSE(r(\rho,\theta), r(\rho+\Delta\rho,\theta)) \quad (7)$$

The displacement $\Delta\rho$ shows how similar the generated images are compared to the cell pattern distribution. To make the calculation simpler, we took $\theta = 0$, $\frac{\pi}{2}$, $\pi$, and $\frac{3}{2}\pi$. Spatial displacement values for SSIM is calculated in the same fashion.

##### 1.3.2. Relative Standard Deviation of Gaussian Noise

We add Gaussian noise with relative standard deviation $\sigma_i$ to the image intensity of a real image $r$, and annotate it as $G(r, 2\sigma_i)$. We then compare the real image to $G$. The smallest $\sigma_i$ that makes the following equation true quantifies the difference between the synthesized and real images compared to added Gaussian noise.

$$MSE(r,g) \leq MSE(r, G(r, 2\sigma_i)) \quad (8)$$

### 2. Supplemental Video Captions

*Supporting Video 1:Aggregate Segmentation*

The aggregate segmentation result of phase-contrast and fluorescent images from 360 min to 1440 min on the test set. Frames prior to 360 min are not included, because the aggregates are not well-formed during that period; the period varies slightly in different biological replicates.

*Supporting Video 2: Model Performance*

Comparison of the real images and images synthesized by pix2pixHD and pix2pixHD-HE in one of the training sets.

### 3. Additional Supplemental Figures

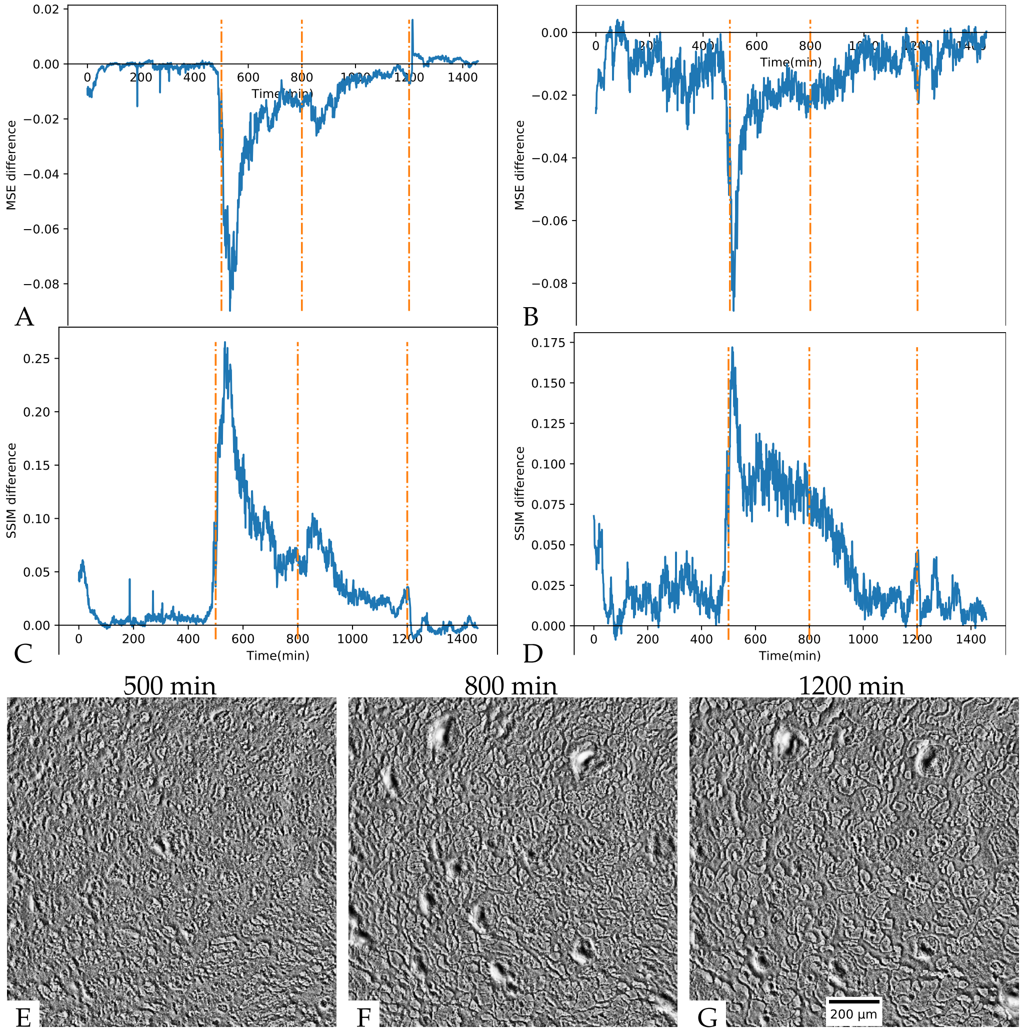

**Figure S4.** Performance evaluation across a training movie. The orange dashed lines show the time points corresponding to E,F,G. (A, B) Subtracting MSE of pix2pixHD from MSE of pix2pixHD-HE in a 24-hour movie; (A) normalized image; (B) Histogram equalized image; (C, D) Subtracting SSIM of pix2pixHD from SSIM of pix2pixHD-HE in a 24-hour movie, (A) normalized image; (B) Histogram equalized image; (E, F, G) phase-contrast image at the 500, 800, 1200 minutes in the movie.

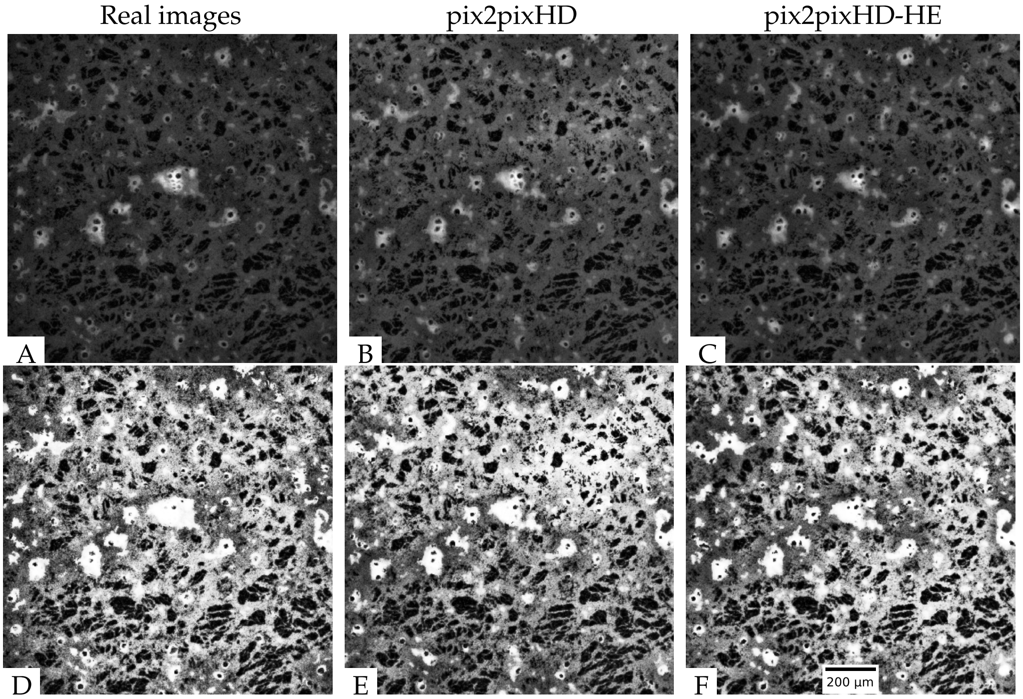

**Figure S5.** Model performance at the 500 min: (A) Ground truth tdTomato fluorescent image; (B) Synthesized fluorescent image (pix2pixHD); (C) Synthesized fluorescent image (pix2pixHD-HE); (D) Histogram equalized ground truth tdTomato fluorescent image; (E) Histogram equalized synthesized fluorescent image (pix2pixHD); (F) Histogram equalized synthesized fluorescent image (pix2pixHD-HE).

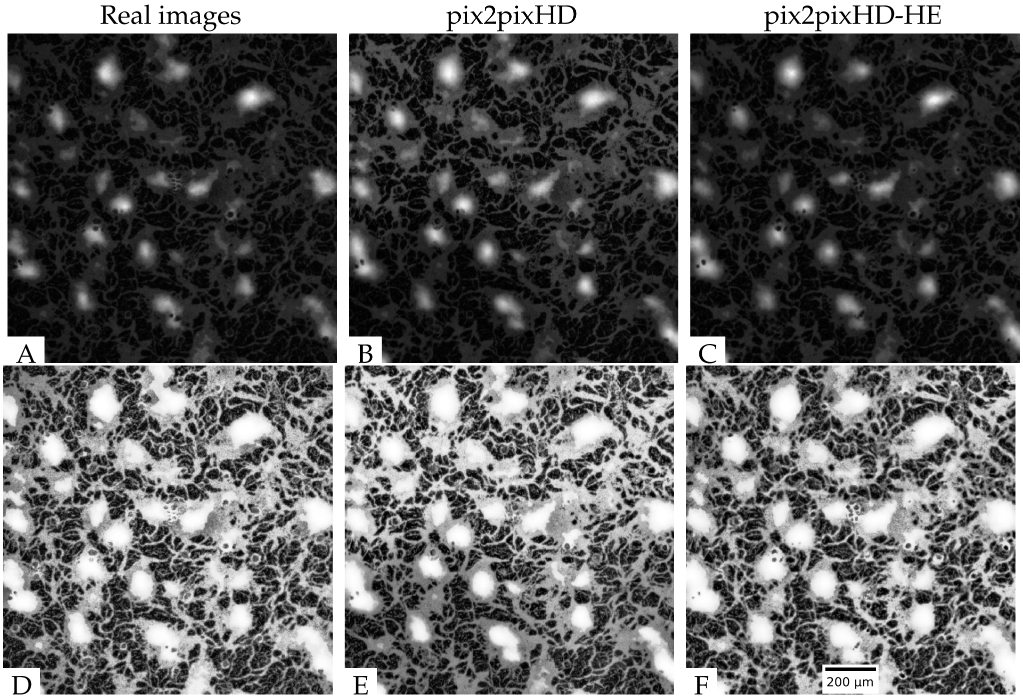

**Figure S6.** Model performance at the 800 min: (A) Ground truth tdTomato fluorescent image; (B) Synthesized fluorescent image (pix2pixHD); (C) Synthesized fluorescent image (pix2pixHD-HE); (D) Histogram equalized ground truth tdTomato fluorescent image; (E) Histogram equalized synthesized fluorescent image (pix2pixHD); (F) Histogram equalized synthesized fluorescent image (pix2pixHD-HE).

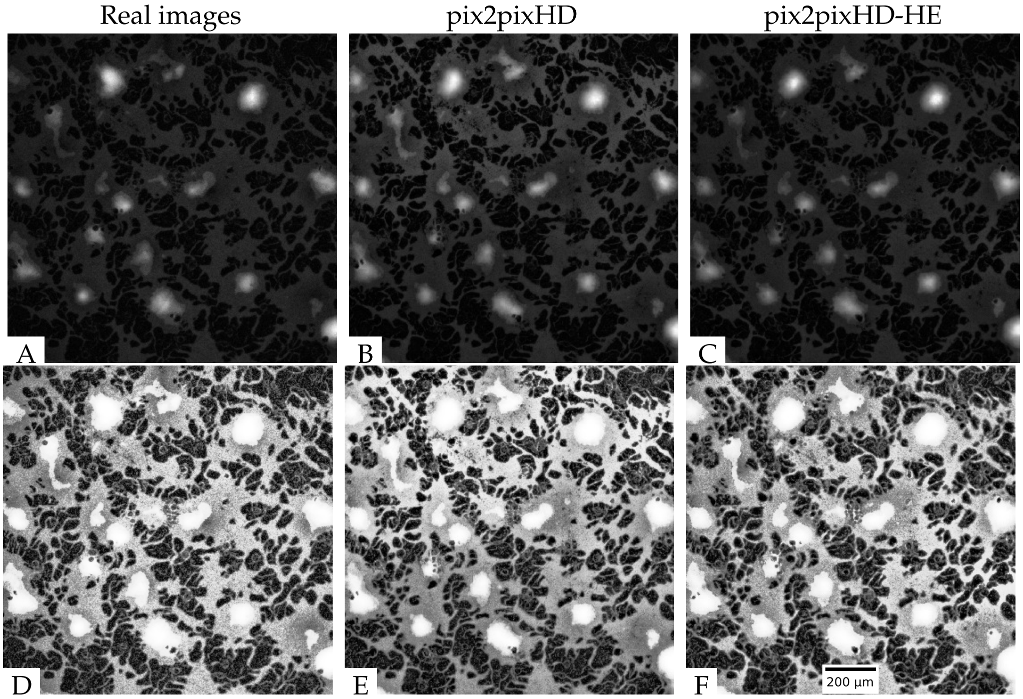

**Figure S7.** Model performance at the 1200 min: (A) Ground truth tdTomato fluorescent image; (B) Synthesized fluorescent image (pix2pixHD); (C) Synthesized fluorescent image (pix2pixHD-HE); (D) Histogram equalized ground truth tdTomato fluorescent image; (E) Histogram equalized synthesized fluorescent image (pix2pixHD); (F) Histogram equalized synthesized fluorescent image (pix2pixHD-HE).

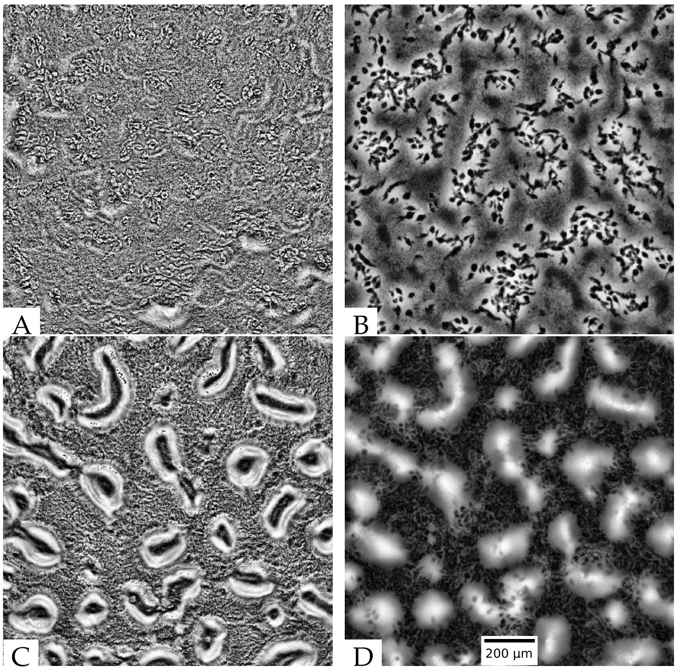

**Figure S8.** Images at the 0 and 800 minutes with cell density $1 \times 10^{10}$ cells/mL: (A) phase-contrast image at the 0 minute; (B) fluorescent image at the 0 minute (the black holes are the dead cells); (C) phase-contrast image at the 800 min; (D) fluorescent image at the 800 min.

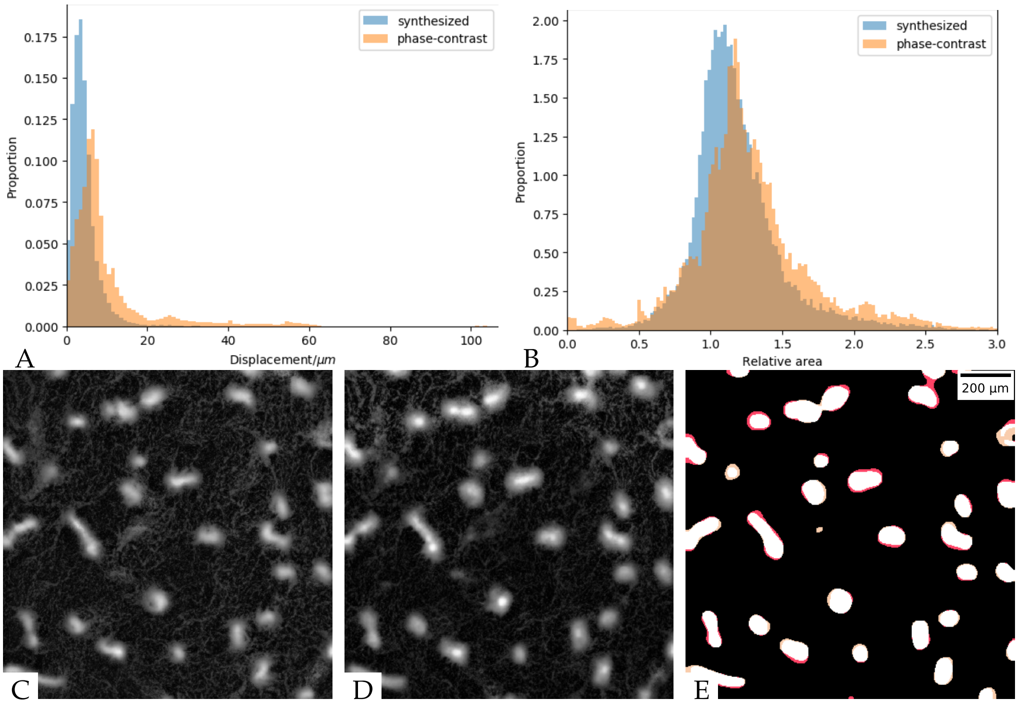

**Figure S9.** Aggregate segmentation for images of $5 \times 10^9$ cells/mL cell density: (A) The distribution of distances between centroid positions. Blue: real and synthesized fluorescent images; orange: phase-contrast and real fluorescent images; (B) The distribution of area ratios for the matched aggregate pairs. Blue: real and synthesized fluorescent images; orange: phase-contrast and real fluorescent images; (C) Fluorescent image at the 800 min; (D) Synthesized image at the 800 min; (E) Aggregate segmentation result at the 800 min. white: true positive, black: true negative, yellow: false positive, red: false negative.

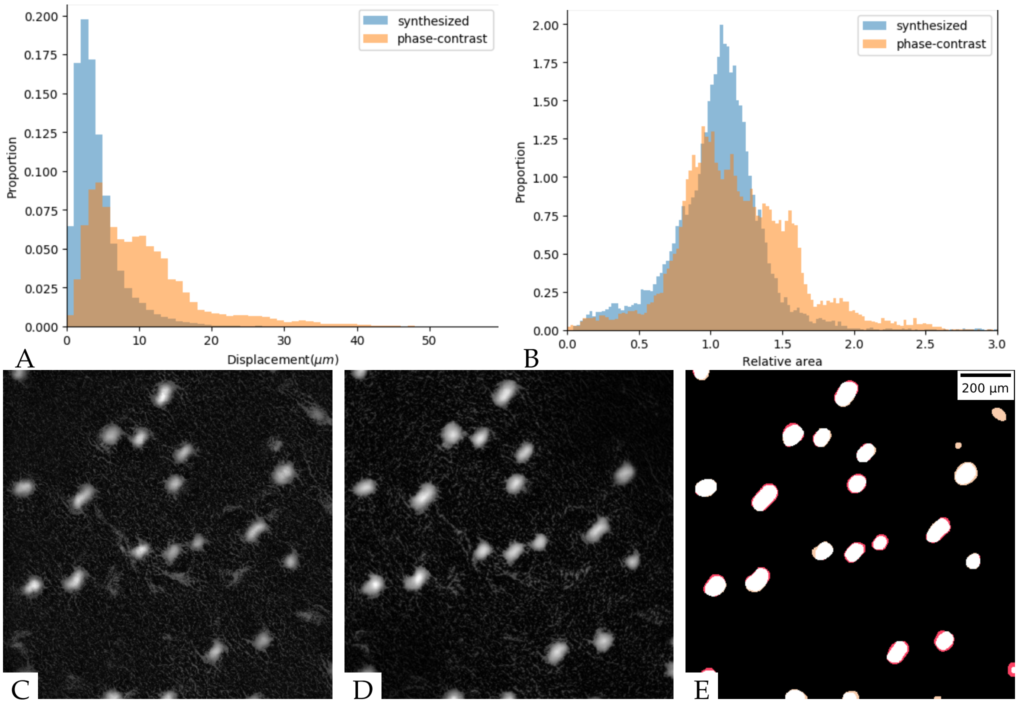

**Figure S10.** Aggregate segmentation for images of $2.5 \times 10^9$ cells/mL cell density: (A) The distribution of distances between centroid positions. Blue: real and synthesized fluorescent images; orange: phase-contrast and real fluorescent images; (B) The distribution of area ratios for the matched aggregate pairs. Blue: real and synthesized fluorescent images; orange: phase-contrast and real fluorescent images; (C) Fluorescent image at the 800th minute; (D) Synthesized image at the 800th minute; (E) Aggregate segmentation result at the 800th minute. white: true positive, black: true negative, yellow: false positive, red: false negative.

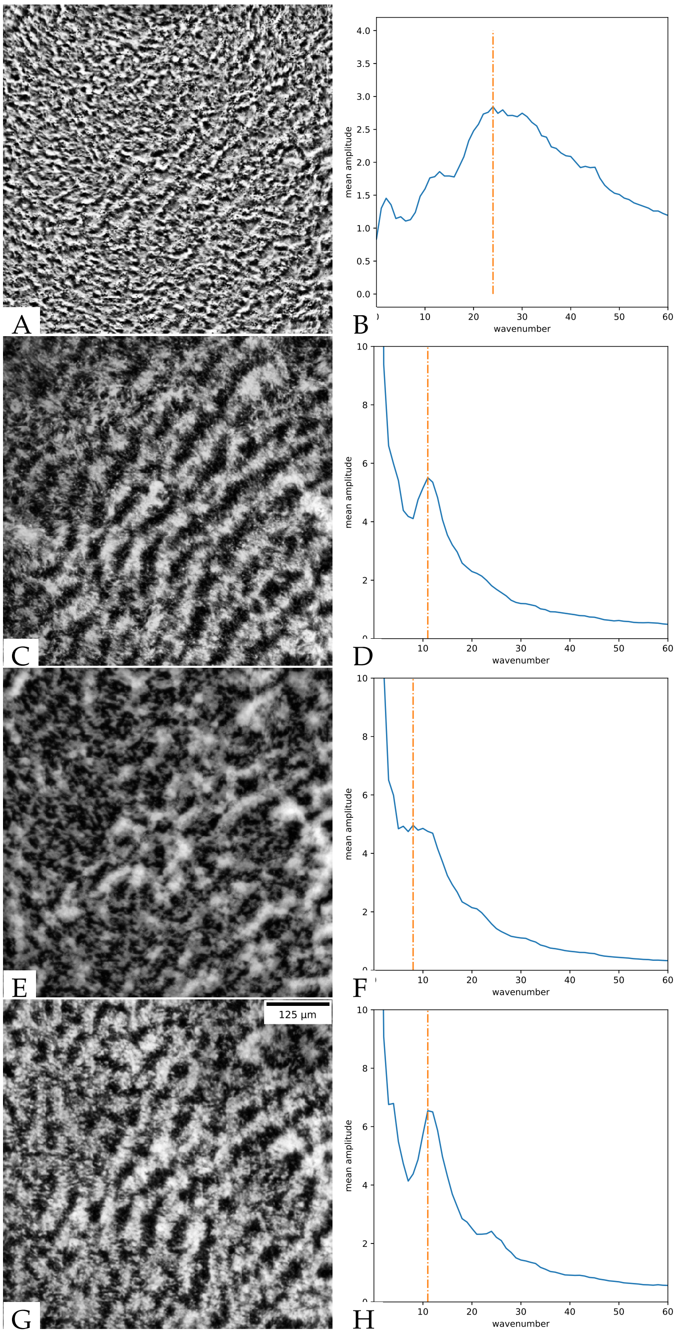

**Figure S11.** The cropped 504 × 504 rippling images and the corresponding mean amplitudes in Fourier space. The wavenumber $k_1$ is shown as an orange dotted line. (A, B) phase-contrast rippling image and the corresponding mean amplitude along the radius, $k_1 = 24, \lambda_k \sim 21\mu m$; (C, D) fluorescent rippling image and the corresponding mean amplitude along the radius, $k_1 = 11, \lambda_k \sim 46\mu m$; (E, F) synthesized fluorescent rippling image before training and the corresponding mean amplitude along the radius, $k_1 = 8, \lambda_k \sim 63\mu m$; (G, H) synthesized fluorescent rippling image after training and the corresponding mean amplitude along the radius, $k_1 = 11, \lambda_k \sim 46\mu m$.

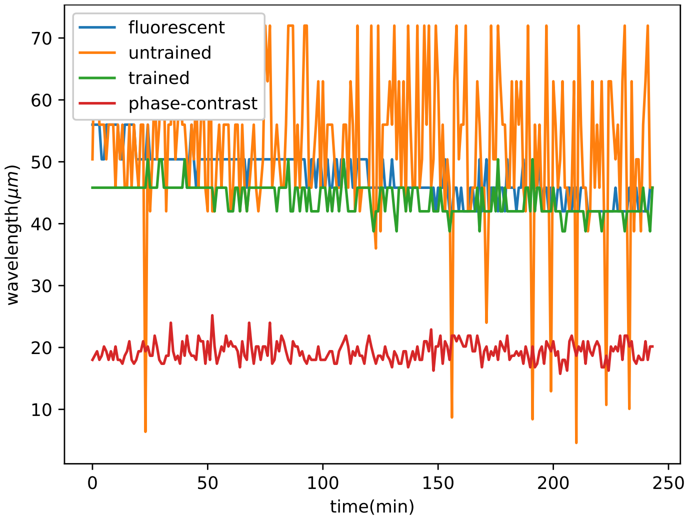

**Figure S12.** The calculated wavelength in a movie. Untrained is the wavelength calculated from the model that was only trained on the aggregation model withour training on ripple patters.Trained is the wavelength calculated from the model further trained on ripple patterns.
